# Learning Orientations: a Discrete Geometry Model

**DOI:** 10.1101/2021.08.08.455577

**Authors:** Yuri Dabaghian

## Abstract

In the mammalian brain, many neuronal ensembles are involved in representing spatial structure of the environment. In particular, there exist cells that encode the animal’s location and cells that encode head direction. A number of studies have addressed properties of the spatial maps produced by these two populations of neurons, mainly by establishing correlations between their spiking parameters and geometric characteristics of the animal’s environments. The question remains however, how the brain may intrinsically combine the direction and the location information into a unified spatial framework that enables animals’ orientation. Below we propose a model of such a framework, using ideas and constructs from algebraic topology and synthetic affine geometry.

## I. INTRODUCTION AND BACKGROUND

**Spatial cognition** in mammals is based on an internalized representation of space—a *cognitive map* ^1^ that emerges from neuronal activity in several regions of the brain [1–3]. The type of information encoded by a specific neuronal population is discovered by establishing correspondences between its spiking parameters and spatial characteristics of the environment. For example, ascribing the *xy*-coordinates to every spike produced by the hippocampal principal neurons according to the animal’s (in the experiments, typically rat’s) position at the moment of spiking, produces distinct clusters, indicating that these neurons, the so-called *place cells*, fire only within specific locations—their respective *place fields* [4, 5]. The layout of the place fields in a spatial domain ε—the place field map *M*_*ε*_ (Fig. 1A)—thus defines the temporal order of the place cells’ spiking activity during the animal’s navigation, which is a key determinant of the cognitive map’s structure. Hence, tagging the spikes with the location information can be viewed as a mapping from a cognitive map 𝒞 into the navigated space,

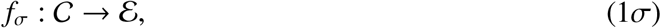

referred to as *spatial mapping* in [6]. Similarly, tagging the spikes produced by certain neurons in the postsubiculum (and in few other brain regions [7, 8]) with the rat’s head direction angle *φ* produces clusters in the space of planar directions—the circle *S* ^1^, thus defining a mapping

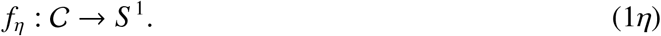

**FIG. 1:**
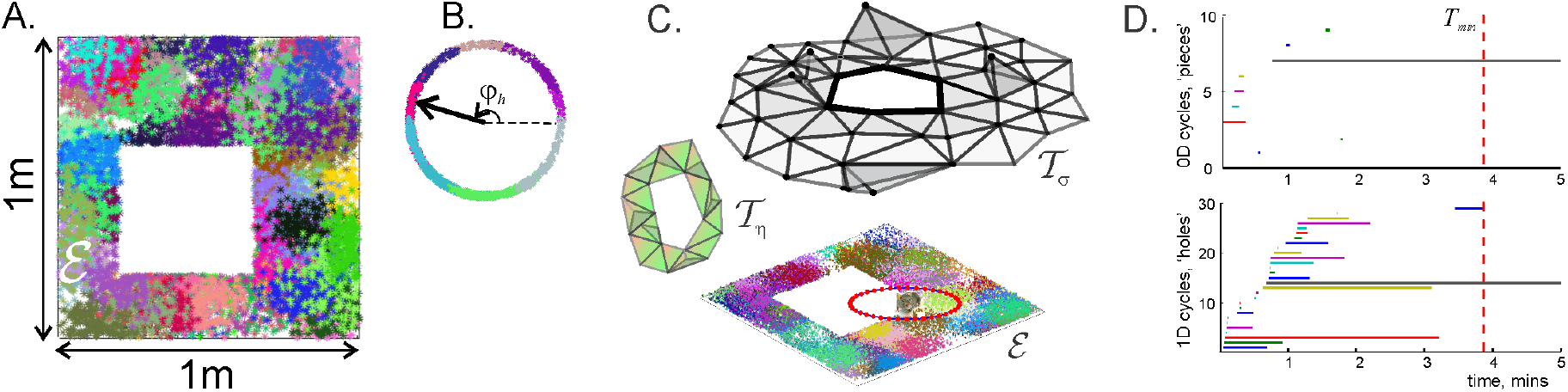
Basic topological constructions. **A**. Simulated place field map *M* _*ε*_ with place fields scattered randomly in a 1 × 1 m square environment E with a square hole in the middle. Clusters of dots of a particular color represent individual place fields. **B**. A head direction field map 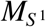 covers the space of directions, *S* ^1^. Clusters of colored dots mark specific head direction fields *υ*_*h*_, centered each at its preferred angle *φ*_*h*_. **C**. The net pool of place cell coactivities is represented by the coactivity complex 𝒯 _*σ*_(*t*) (top right), which provides a developing topological representation of the environment *ε* (bottom). The head direction cells map a circular space of directions *S* ^1^ (shown as a ring around the rat). The net pool of head direction cell activities is schematically represented the coactivity complex 𝒯_*σ*_(*t*). **D**. The timelines of the separate pieces (top panel) and holes (bottom panel) in the complex 𝒯 _*σ*_(*t*) are shown as horizontal bars. At the onset of the navigation, 𝒯 _*σ*_(*t*) contains many spurious topological defects that disappear after a certain “learning period” 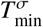, leaving behind a few persistent loops that define the topological shape of 𝒯 _*σ*_(*t*) [38]. Similar behavior is exhibited by the head direction coactivity complex 𝒯_*η*_(*t*).

The angular domains in which specific *head direction cells* become active can be viewed as *head direction fields* in *S* ^1^, similar to the hippocampal place fields in the navigated space. The corresponding *head direction map*, 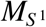, determines the order in which the head direction cells spike during the rat’s movements (Fig. 1B, [9, 10]).

The preferred angular domains depend weakly, if at all, on the rat’s position, just as place fields are overall decoupled from the head or body orientation (see however [11, 12]). Thus, the following discussion will be based on the assumption that both cell populations contribute to an *allocentric* representation of the ambient space: the place cells encode a topological map of locations [13–18], whereas head direction cells augment it with angular information [19–22].

### Topological model

The physiological and the computational mechanisms by which a cognitive map comes into existence remain vague [23–25]. However, certain insights into its structure can be obtained through geometric and topological constructions. For example, a place field map *M*_*ε*_ can be viewed as a *cover* of the navigated environment *ε* by the place fields *υ*_*i*_,

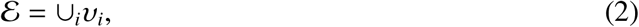

and used to link the topology of *ε* to the topological structure of the cognitive 𝒞 map. Indeed, according to the Alexandrov-Čech theorem, if every nonempty set of overlapping place fields, 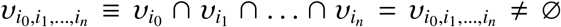, is represented by an abstract simplex, 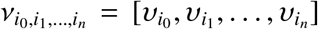, then the homologies of the resulting simplicial complex 𝒩_*σ*_—the *nerve* of the map *M*_*ε*_—match the homologies of the underlying space *H*_*_(𝒩_*σ*_) = *H*_*_(*ε*), provided that all the overlaps 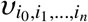 are contractible. This implies that 𝒩_*σ*_ and εhave the same *topological shape*— same number of connectivity components, holes, cavities, tunnels, etc. [26]. The same line of arguments allows relating the head direction map with the topology of the space of directions *S* ^1^ (Fig.1C).

It must be emphasized however, that reasoning in terms of place and head direction fields may not capture the brain’s intrinsic principles of processing spiking information, e.g., explain how either the location or the direction signals contribute to animal’s spatial awareness, because the experimentally constructed firing fields are nothing but artificial constructions used to in interpret and visualize spiking data [29, 30]. Addressing the brain’s intrinsic space representation mechanisms requires carrying the analyses directly in terms of spike times, without invoking auxiliary correlates between neuronal activity and the observed environmental features.

Fortunately, the approach motivated by the nerve theorem can be easily transferred into a “spiking” format. Indeed, one can view a combination of the coactive place cells—a *cell assembly* [31–33]—as an abstract *coactivity simplex*,

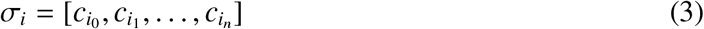

that activates when the rat crosses its *simplex field* 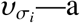 domain where all the cells *c*_*i*_ ∈ *σ*_*i*_ are coactive [34]. By construction, this domain is defined by the overlap of the corresponding place fields 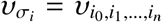, and may hence be viewed as the the projection of *σ*_*i*_s into *ε*under the mapping (1*σ*). Note that if two coactivity simplexes overlap, their respective fields also overlap, 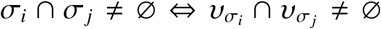. Thus, if a cell *c*_*i*_ is shared by a set *U*_*i*_ of simplexes, *U*_*i*_ = {*σ* : *σ* ∩ *c*_*i*_ ≠ ∅}, then its place field is formed by the union of the corresponding *σ*-fields,

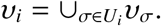

If a simplex *σ*_*i*_ first appears at the moment *t*_*i*_, then the net pool of neuronal activities produced by the time *t* gives rise to a time-developing simplicial *coactivity complex*

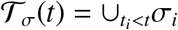

inflates (𝒯_*σ*_(*t*) ⊆ 𝒯_*σ*_(*t*′) for *t < t* ′), and eventually *saturates*, converging to the nerve complex’s structure, i.e., 𝒯_*σ*_(*t*) ≈ 𝒩_*σ*_, for 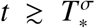. Analyses based on simulating rat’s moving through that randomly scattered place fields show that, e.g., for a small environment *ε* illustrated on Fig. 1A, the rate of new simplexes’ appearance slacks in about 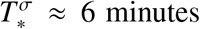 [6], which provides an estimate for the time required to map *ε*.

The topological dynamics of 𝒯_*σ*_(*t*) can be described using Persistent Homology theory [35– 37], which allows identifying the ongoing shape of 𝒯_*σ*_(*t*) based on the times of its simplexes’ first appearance. Typically, 𝒯_*σ*_(*t*) starts off with numerous topological defects that tend to disappear as the information provided by the spiking place cells accumulates (see [38–41] and Fig. 1D). Hence the minimal period 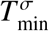 required to recover the “physical” homologies *H*_*_(*ε*) provides an estimate for the time necessary to learn topological connectivity of the environment, which, for the case illustrated on Fig. 1A, is about 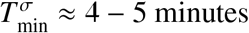 [38–43].

Importantly, the coactivity complex may be used not only as a tool for estimating learning timescales, but also as a schematic representation of the cognitive map’s developing structure, providing a context for interpreting the ongoing neuronal activity. Indeed, a consecutive sequence of *σ*-fields visited by the rat,

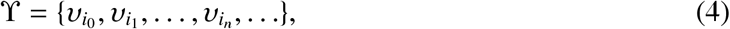

captures the shape of the underlying physical trajectory *s* ⊂ ϒ [44–48]. The corresponding chain of the place cell assemblies ignited in the hippocampal network is represented by the *simplicial path*

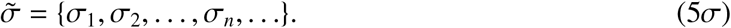

The fact that this information allows interpreting certain cognitive phenomena [49–51] suggests that the animal’s movements are faithfully monitored by neuronal activity, i.e., that in sufficiently well-developed complexes (e.g., for 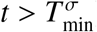) simplicial paths capture the shapes of the underlying trajectories [44–47]. For the referencing convenience, this assumption is formulated as two model requirements:

**R1. Actuality**. *At any moment of time, there exists an active assembly σ that represents the animal’s current location*.

**R2. Specificity**. *Different place cell assemblies represent different domains in ε, i*.*e*., *σ-simplexes serve as unique indexes of the animal’s location in a given map* 𝒞.

An implication of these requirements is that the simplex fields cover the explored surfaces (2) and that if the consecutive simplexes in (5*σ*) are *adjacent*, i.e., no simplexes ignite between *σ*_*i*_ and *σ*_*i*+1_ (schematically denoted below as 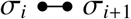) then the corresponding *σ*-fields are adjacent or overlap.

### Head orientation map

Using the same line of arguments, one can deduce the topology of the space of directions by building a dynamic *head direction coactivity complex* 𝒯_*η*_(*t*) from the simplexes

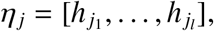

which designate the assemblies of head direction cells *h*_*j*1_, *h*_*j*2_, …, *h*_*jl*_. If a simplex *η*_*j*_ first activates at the moment *t* _*j*_, then

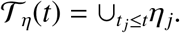

As the complex 𝒯_*η*_(*t*) develops, it forms a stage for representing the head direction cell spiking structure: in full analogy with (5*σ*), traversing a physical trajectory *s*(*t*) induces a sequence of active *η*-simplexes, or a head direction simplicial path

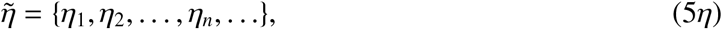

in which different *η*-simplexes represent distinct directions, at all locations. As spiking information accumulates, the topological structure of 𝒯_*η*_(*t*) converges to the structure of nerve complex 𝒩_*η*_ induced by the head direction fields’ cover of *S* ^1^—every *η*-simplex projects into its respective head direction field *υ*_*η*_ under the mapping (1*η*). Simulations demonstrate that in the environment shown on Fig. 1A, a typical coactivity complex 𝒯_*η*_(*t*) saturates in about 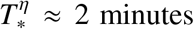, while the persistent homologies of 𝒯_*η*_(*t*) filtered according to the times of *η* –simplexes’ first appearances reveal the circular topology of the space of directions in about 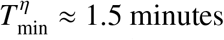.

From the biological point of view however, these results do not provide an estimate for *orientation learning time*: by itself, 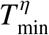 may be viewed as the time required to learn head directions at a particular location, in every environment, whereas learning to orient in *ε* implies knowing directions at every location and an ability to link orientations across locations. The latter is a much more extensive task, which, as it will be argued below, requires additional specifications and interpretations.

The following discussion is dedicated to constructing phenomenological models of orientation learning using algebraic topology and synthetic geometry approaches. In Section II, we construct and test a direct generalization of the topological model, similar to the one used in [38–41] and demonstrate that it fails to produce biologically viable predictions for the learning period. In Section III, the topological approach is qualitatively generalized using an alternative scope of ideas inspired by synthetic geometry. In Section IV, it is demonstrated that the resulting framework allows incorporating additional neurophysiological mechanisms and acquiring the topological connectivity of the environment in a biologically viable time, thus revealing a new level of organization of the cognitive map, as discussed in Section V.

## II. TOPOLOGICAL MODEL OF ORIENTATION LEARNING

### Orientation coactivity complex

The model requirements **R1** and **R2**, applied to both hippocampal and head direction activity, imply that the animal’s location and orientation are represented, at any moment of time, by an active *σ*-simplex and an active *η*-simplex. Thus, the net pattern of activity in the hippocampal and in the head direction networks defines a (*σ, η*) pair—a single *oriented*, or *pose* simplex

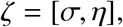

(the latter term is borrowed from robotics [22, 52, 53]). Restricting a *ζ*-simplex to its maximal subsimplexes spanned, respectively, by the place- or the head direction cells defines the projections into its positional and directional components,

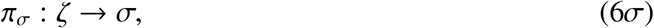

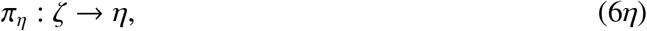

which permits terminology such as “*ζ* is located at *σ*,” “*ζ* is directed toward *η*,” “a location *σ* is directed by *η*,” “*η* is applied at *σ*,” etc. Thus, one may refer to the *σ*-simplexes as to *locations* and to the *η*-simplexes as to *directions*, implying, depending on the context, either the items encoded in the cognitive map, or the *σ/η*-fields, or both.

As in the previously discussed cases, the collection of pose simplexes produced up to a moment *t* forms an *orientation coactivity complex* 𝒯 _*ζ*_(*t*) that schematically represents the net pool of conjunctive patterns generated by the place- and the head direction cells accumulated since the onset of the navigation. In particular, the combinations of cells ignited along a physical path *s* induces an *oriented simplicial path*

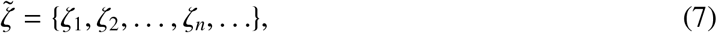

which runs through 𝒯_*ζ*_(*t*). The transitions from a given active pose simplex, *ζ*_*i*_, to the next, *ζ*_*i*+1_, occur at discrete moments *t*_1_, *t*_2_, …, *t*_*n*_, …, when either the *σ*- or the *η*-component of *ζ*_*i*_ deactivates and the corresponding component of *ζ*_*i*+1_ ignites. Thus, the simplicial paths 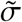 and 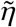 can be pro- duced from the oriented path (7) using (6). In contrast with (5*σ*) and (5*η*), the *σ*- and *η*-simplexes in such paths are indexed uniformly, according to the indexes of (7),

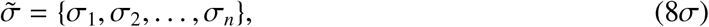

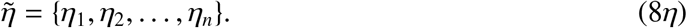

The adjacent simplexes in either (8*σ*) or in (8*η*) (but not in both of them simultaneously) may coincide, e.g., the location *σ*_*i*_ may remain the same during several timesteps, while the *η*-activity changes, or vice versa.

Since the rat can potentially run in any direction at any location (unless stopped by an obstacle), there are no *a priori* restrictions on the order of the place cell and the head direction cells spiking activity. This observation is formalized by another model requirement:

**R3. Independence**. *A given head direction cell assembly η may become coactive with any place cell assembly σ and vice versa, with independent σ- and η-spiking parameters*.

In model’s terms, this implies that the development of the coactivity complex 𝒯 _*η*_(*t*) and its ultimate saturated structure 𝒩 _*η*_ is the same at any location *σ*, and vice versa, the *saturated* structure of 𝒯 _*σ*_(*t*) is independent from *η*-activity.

However, since the activities in the hippocampal and in the head direction cell networks represent complementary aspects of the same movements, certain characteristics of the simplicial paths (8) are coupled. Specifically, in light of **R1**–**R2**, a connected physical trajectory *s* should induce a connected *σ*-path in the place cell complex 𝒯 _*σ*_, together with a connected *η*-path in the head direction complex 𝒯 _*η*_. Similarly, a *looping* trajectory should induce periodic sequences of simplexes,

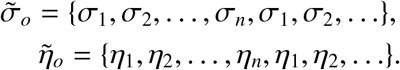

In other words, making a loop in physical space *ε* should induced *σ*- and *η*-loops. Thus, without referencing the physical trajectory, the model requires

**R4. Topological consistency**. *The simplicial paths (8) should be connected and a simple periodic σ-path should induce a simple periodic η-path and vice versa*.

### Orientation learning

As discussed above, getting rid of the topological defects in 𝒯_*σ*_(*t*) and in 𝒯_*η*_(*t*) allows faithful topological classification of physical routes in terms of the neuronal (co)activity. Thus, the “topological maturation” of these complexes can be viewed as a schematic representation of the learning process. The concept of oriented simplicial paths embedded into the orientation coactivity complex 𝒯_*ζ*_(*t*) allows a similar interpretation of the spatial orientation learning—as acquiring an ability to distinguish between qualitatively disparate moving sequences. Indeed, knowing how to orient in a given space, viewed as a cognitive ability to reach desired places from different directions via a suitable selection of intermediate locations and turns, may be interpreted mathematically as an ability to classify trajectories using topologically inequivalent classes of oriented simplicial paths (8). From an algebraic-topological perspective, this may be possible after the orientation complex 𝒯_*ζ*_(*t*) acquires its correct topology.

To establish the latter, note that the complex 𝒯_*ζ*_(*t*) has the same nature as 𝒯_*σ*_(*t*) and 𝒯_*η*_(*t*)—it is an emerging temporal representation of a nerve complex, induced from a cover of a certain *orientation space* 𝒪that combines *ε* and *S* ^1^. Since the preferred angles of the head direction cells remain the same at all locations [7], the space of directions *S* ^1^ represented by these cells does not “twist” as the rat moves across *ε*, which implies that the orientation space has a direct product structure 𝒪= *ε* ×*S* ^1^ [26]. One may thus combine (1*σ*) and (1*η*) to construct a *joint spatial mapping, f*_*ζ*_ = (*f*_*σ*_, *f*_*η*_), that associates instances of simultaneous activity of place- and head direction cell groups with domains in the orientation space,

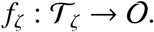

For example, if a given place cell *c* maps into a field *υ* = *f*_*σ*_(*c*), then the coactivity of a pair *ζ* = [*c, h*] (the smallest possible combined coactivity) can be mapped into 𝒪 by shifting *υ* along the corresponding fiber *S* ^1^ according to the angle field of the head direction component of *ζ, φ* = *f*_*η*_(*h*) (Fig. 2A). The resulting *orientation fields*, 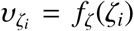, form a cover of the orientation space,

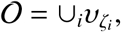

**FIG. 2:**
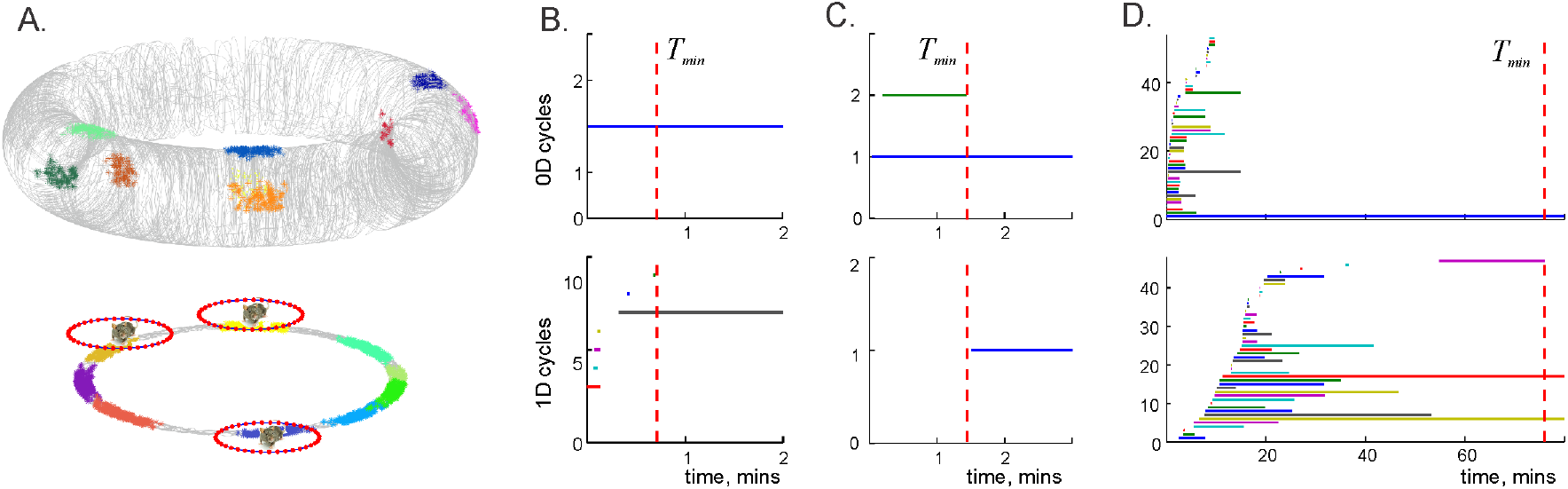
Topological leaning dynamics. **A**. The orientation space for a rat moving on a circular runway (bottom) is a topological torus (top), 𝒪 = *T* ^2^. Clusters of colored dots show examples of the simplex fields 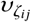 of the basic coactivity combinations *ζ*_*i j*_ = [*c*_*i*_, *h*_*j*_]. The timelines of the topological loops in the place cell complex 𝒯_*σ*_(*t*) (panel **B**, horizontal bars) and in the head direction cell complex 𝒯_*η*_(*t*) (panel **C**) disappear in under a minute; a stable loop in 0*D* and a stable loop in 1*D* in each case indicate that both the runway and the direction space are topological circles. **D**. The orientation coactivity complex 𝒯_*ζ*_ (*t*) contains many topological defects that take over an hour to disappear: the spurious 0*D* loops contract in about 17 minutes (top panel), and the spurious 1*D* loops persist for 80 minutes. For an “open field” environment (Fig. 1A) a similar topological learning process takes hours. These estimates exceed the experimental learning timescales, suggesting that the physiological orientation learning involves additional mechanisms.

whose nerve 𝒩_*ζ*_ is reproduced by the temporal orientation complex 𝒯_*ζ*_(*t*).

Just as the *σ*- and *η*-simplexes, each pose simplex *ζ*_*k*_ has a well-defined appearance time, *t*_*k*_, due to which the orientation complex is *time-filtered*, 𝒯_*ζ*_(*t*) ⊆ 𝒯_*ζ*_(*t*′) for *t < t*′. Applying Persistent Homology techniques [35, 36], one can compare topological shape defined by the time-dependent Betti numbers of 𝒯_*ζ*_(*t*) with the shape of the orientation space = 𝒪 × *ε S* ^1^, and thus quantify the orientation learning process.

As an illustration of this approach, we simulated the rat’s movements on a circular runway, for which the total representing space for the combined place cell and head direction cell coactivity forms a 2*D* torus (Fig. 2A). To simplify modeling, we used movement direction as a proxy for the head direction, although physiologically these parameters not identical [54–58]. Computations show that the correct topological shapes of the place- and the head direction complexes emerge in about 1 minute (Fig. 2B,C), while the transient topological defects in 𝒯_*ζ*_(*t*) disappear in about 80 minutes (Fig. 2D)—a surprisingly large value that exceeds behavioral outcomes by an order of magnitude [42, 43]. Even assuming that biological learning may involve only a partial reconstruction of the orientation space’s topology, the persistence diagrams shown on Fig. 2D indicate that 𝒯_*ζ*_(*t*) is not only riddled with holes for over 40 minutes, but also that it remains disconnected (*b*_0_(𝒯_*ζ*_) *>* 1) for up until 17 minutes, i.e., according to the model, the animal should not be able to acquire a connected map of a simple annulus after completing multiple lapses across it.

**Biological implications** of mismatches between the experimental and the modeled estimates of learning timescales are intriguing: since the model’s quantifications are based on “topological accounting” of place- and head direction cell coactivities induced by the animal’s movements, the root of the problem seems to lay not in the mathematical side of the model, but in the biological assumptions that underlie the computations. Specifically, the overly long learning periods suggest that building a spatial map from the movement-triggered neuronal coactivity alone does not take into account certain principal components of the learning mechanism. In other words, the fact that the animal seems to produce correct representations of the environment much faster than it would be possible from the influx of navigation-triggered data, implies that the brain can bypass the necessity to discover every bit of information empirically, i.e., that building a cognitive map may be accelerated by “generating information from within,” via autonomous network dynamics. Physiologically, this conclusion is not surprising: the phenomena associated with spatial information processing through endogenous hippocampal activity are well known. Many experiments have demonstrated that the animal can *replay* place cells during the quiescent states [18, 59, 60] or sleep [61, 62], in the order in which they have fired during preceding active exploration of the environment, or *preplay* place cells in sequences that represent future trajectories [49–51]. These phenomena are commonly viewed as manifestations of the animal’s “mental explorations” of its cognitive map, which help acquiring, sustaining and retrieving memories [63, 64].

However, one would expect a different functional impact of replays and preplays on spatial learning. Since replays represent past experiences, they cannot accelerate acquisition of new spatial information—in the model’s terms, reactivation of simplexes that are already included into the coactivity complex cannot alter its shape (in absence of synaptic and structural instabilities [65–68]). In contrast, preplaying place cell combinations that have not yet been triggered by previous physical moves may speed up the learning process. Yet, from the modeling perspective, it is *a priori* unclear which specific trajectories may be preplayed by the brain, or, in computational terms, which specific simplicial trajectories should be “injected” into the simulated cognitive map 𝒯_*ζ*_(*t*) to simulate the preplays that may accelerate learning. Experiments suggest that “natural” connections between locations are the straight runs [20, 21, 49]; however, implementing such runs would require a certain “geometrization” of the topological model. In the following, we propose a geometric implementation of preplays that help to expedite learning process and open new perspectives modeling spatial representations.

## III. GEOMETRIC MODEL OF ORIENTATION LEARNING

**The motivation** for an alternative approach comes from the observation that combining inputs from the place and the head direction cells offers a possibility of establishing different arrangements of the locations in the hippocampal map. Indeed, common interpretations of the head direction cells’ functions suggest that the rat’s movements guided by a fixed head direction activity trace approximately straight paths, whereas shifts in *η*-activity indicate curved segments of the trajectory, turns, etc. [7, 22, 69]. This implies that the brain may use the head direction cells’ outputs to represent shapes of the paths encoded by the place cells, to align their segments, identify collinearities, their incidences, parallelness, etc.

To address these structures and their properties we will use the following definitions:

**D1**. Simplexes *σ*_1_ and *σ*_2_ are *aligned in η-direction*, if they may ignite during an uninterrupted activity of a fixed *η*-simplex. In formal notations, (*σ*_1_, *σ*_2_) ◁ *η*.

**D2**. Two *η*-aligned simplexes are *η*-*adjacent*, 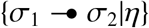, if the ignition of *σ*_2_ follows immediately the ignition of *σ*_1_, with no other cell groups igniting in-between.

**D3**. An ordered sequence of *σ*-simplexes forms an *η-oriented alignment* if each pair of consecutive simplexes, (*σ*_*i*_, *σ*_*i*+1_) in (8*σ*), is *η*-adjacent, i.e., if the oriented path,

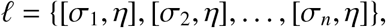

never changes direction. The notation

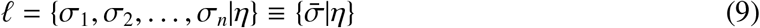

highlights the set of *collinear* locations 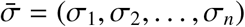 and the *η*-simplex that orients it. The bar in 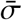 is used to distinguish an alignment from a generic simplicial path 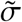.

**D4**. An alignment _1_ *augments* an alignment _2_, if both _1_ and _2_ can be guided by an uninterrupted *η*-activity 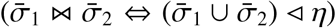. Conversely, a proper subset 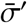 of an aligned set 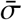 forms its proper *subalignment* 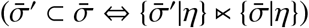.

**D5**. Two alignments 𝓁_1_ and 𝓁_2_ *overlap*, if they share a location 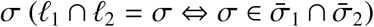.

**D6**. A location *σ*′lays *outside* of an *η*-alignment 𝓁_*η*_, if it aligns with any *σ* from 𝓁_*η*_ along a direction different from *η* (*σ*′ ∉ 𝓁_*η*_ ⇔ ∃*σ* ∈ 𝓁_*η*_′ *η* ≠*η* : (*σ, σ′*) ◁ *η*′).

**D7**. Two alignments are *parallel*, if they are directed by the same or opposite *η*-activity, without augmenting each other, i.e., if one ± *η*-alignment contains a location outside of the other one, (𝓁 _1_ II 𝓁_2_ ⇔ *η*_1_ = ± *η*_2_, and 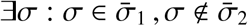, where *f*_*η*_(−*η*) ≊ *π* + *f*_*η*_(*η*)).

**D8**. A *yaw* is an oriented path in which a sequence of *η*-simplexes ignites at a fixed location *σ*,

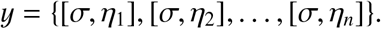

Thus, yaws may be viewed as structural opposites of the alignments, which is emphasized by the notation

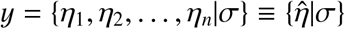

that highlights the range of *η*-simplexes, 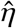, ignited at the *axis of the yaw, σ*.

**D9**. A *clockwise turn* is an oriented path 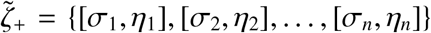 with a growing angular sequence, i.e., the angle *φ*_*i*_ that represents the element *η*_*i*_ is not greater than the next one, *φ*_*i*+1_ ≥ *φ*_*i*_. A *counterclockwise turn* 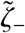 is an oriented path with a decreasing angular sequence, *φ*_*i*+1_ ≤*φ*_*i*_.

The latter definition is due to the observation that *η*-simplexes can be ordered according to the angles they represent, i.e., *η*_*i*_ *< η*_*i*+1_ iff *φ*_*i*_ *< φ*_*i*+1_, which also allows defining the angle between alignments,

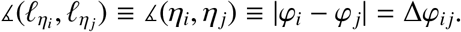

In light of the definitions **D1**–**D9**, the model requirements **R1**–**R4** imply that the entire ensembles of the active place and head direction cells are involved into geometric arrangements, e.g., every location *σ* belongs to an alignment directed by a *η*-simplex, and conversely, every *η*-simplex directs a nonempty alignment through the *σ*-map. The question is, whether this collection of alignments is sufficiently complete to allow self-contained geometric reasoning in terms of collinearities, incidences, parallelisms, etc., i.e., does it form a self-contained geometry?

The standard approach to answering this question is based on verifying a set of axioms, in this case—the axioms of affine geometry, applied to the elements of a suitable set *A* = {*x, y, z*, …} and its select subsets 𝓁_1_, 𝓁_2_, …, ∈ *A*:

**A1**. *Any pair of distinct elements of x* ≠ *y is included into a unique subset*. 𝓁

**A2**. *There exists an element x outside of any given subset* 𝓁, *x* ∉ 𝓁.

**A3**. *For any subset and an element x* ∉ 𝓁., *there exists a unique subset* 𝓁_*x*_ *that includes x, but does not overlap with*. 𝓁

If these axioms (referred to as ***A****-axioms* below) are satisfied, then the subsets 𝓁_1_, 𝓁_2_, …, can be viewed as lines because interrelationships among them and with other elements of *A* reproduce the familiar geometric incidences between lines and points in the Euclidean plane [70–73]. However, the set of geometries established via the **A**-axioms is much broader than its main “motivating example:” the standard planar affine geometry 𝒜_*E*_ is but a specific model implementing the **A**-axioms using the infinite set of infinitesimal points and infinite lines [70–73]. In fact, it is also possible to use the **A**-axioms to establish geometry on finite sets, thus producing finite affine planes. This is important for modeling cognitive maps encoded by the physiological networks that contain finite numbers of neurons and thus may represent finite sets of locations and alignments. Specifically, a possible adaptation of the **A**-axioms using spiking semantics, is the following:

**A1**_*n*_. *Any pair of distinct locations σ*_1_ ≠ *σ*_2_ *belongs to a unique alignment, i*.*e*., *σ*_1_ *and σ*_2_ *may ignite in sequence during the activity of a single η-simplex* (∀*σ*_1_, *σ*_2_, ∃!*η*,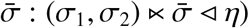).

**A2**_*n*_. *The location-encoding network can activate a group of cells σ to represent a location outside of any given alignment* (∀𝓁, ∃*σ* (∉. 𝓁).

**A3**_*n*_. *For any alignment and a location σ* ∉ 𝓁., *there exists a unique alignment* 𝓁_*σ*_ *parallel to* 𝓁 *that passes through σ* (∀𝓁, *σ∉*, ∃!𝓁_*σ*_ : *σ* ∈ 𝓁_*σ*_, 𝓁 II 𝓁_*σ*_).

Validating these axioms over the net pool of spiking activities produced by the hippocampal and head direction cells would establish a discrete-geometric structure encoded by (*σ, η*)-neuronal activity. However, the requirements imposed by the **A**_*n*_-axioms may not be compatible with physiological mechanisms that operate the corresponding networks, as well as with these networks’ functions. Indeed, the configurations formed by the connections in finite planes are typically non-planar (Fig. 3), whereas physiological computations combining place and head direction cells’ activities appear to enable geometric planning in planar environments [20, 21]. Second, the combinations of locations that form “relational lines” according to the **A**_*n*_-axioms may not have the standard properties of their Euclidean counterparts, e.g., they may include sequences of locations that cannot be consistently mapped into straight Euclidean paths (Fig. 3). In contrast, experiments show that neuronal activity during animals’ movements along straight arrangements of *σ*-fields, as well as their offline preplays/replays [75, 76], dovetail with the definitions **D1**-**D7**. Third, given a large number of the encoded locations (in rats, about 3 × 10^4^ of active place cells in small environments [77]) and a very large set of possible co-active cell combinations [33, 79], a finite set of *η*-simplexes may not suffice to align all pairs of locations in the sense of the definition **D7**.

**FIG. 3:**
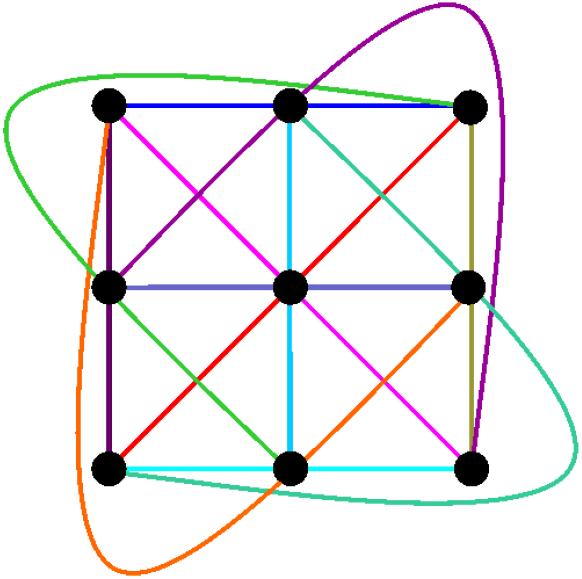
Finite affine plane of the third order, 𝒜_3_, with 9 points (black dots) aligned according to the **A**-axioms in 12 lines (colors) forms a non-planar configuration.

On the other hand, certain key features of finite affine planes, e.g., the necessity of having *k* parallel lines in every direction and a fixed number, *k* + 1, of lines passing through each location [70–73] are reflected in the network. Indeed, the existence of a fixed population of head direction assemblies results in a fixed number *k* ≊ *N*_*η*_*/*2 of distinct alignments passing through any location and the same number of directions running across the cognitive map.

Together, these observations suggest that the brain may combine certain aspects of finite geometries and Euclidean plane discretizations. Rather than trying to recognize the net geometry of the resulting representations from the onset, one may adopt a constructive “bottom up” approach: it may be possible to interpret certain *local* properties of spiking activity as basic geometric relations and then follow how such relations accumulate at larger scales, yielding *global* geometric frameworks. For example, it can be argued that place cell activities can be aligned locally, i.e., that ignitions of a specific head direction assembly can accompany transitions of activity from one place cell assembly to an adjoining one. It is also plausible that, in stable network configurations, the selection of cell groups involved in such transitions is limited or even unambiguous. Also, given the number of place cells (*N*_*c*_ ≈ 3 × 10^5^) and typical assembly sizes (60 − 300 cells) [33], there should be enough place cell combinations to represent a sufficient set of alignments, overlaps between them, etc. [78–80].

Thus, in addition to (mostly topological) requirements **R1**–**R4**, one can consider the following neuro-geometric rules:

**G1**. *Any two adjacent locations align in a unique direction* (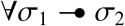: (∀*σ*_1_, *σ*_2_) ◁ *η*).

**G2**. *A location adjacent to a given one may be recruited in any direction* (∀*η, σ*_1_, 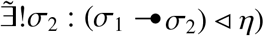).

**G3**. *The location-encoding network can explicitly represent the overlap between any two non-parallel alignments* (∀𝓁_1_, 𝓁_2_, *η*_1_ ≠ *η*_2_, 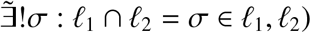).

In contrast with the **A**_*n*_-axioms that aim to establish large-scale properties of a “cognitive” affine plane as a whole, the **G**-rules define local geometric relationships induced by local mechanisms controlling neuronal activity. In particular, **G1** ascertains a possibility of aligning any two adjacent locations, rather than any two locations as required by the **A1**_*n*_. The rule **G2** is complementary: it posits that if an active *η*-combination is selected, then the activity can propagate from a given *σ*_1_ to a specific adjacent *σ*_2_. Lastly, the rule **G3** allows reasoning about the locations, alignments, incidences, etc., assuming that all these elements can be physiologically actualized.

As a first application of the **G**-rules, notice that a *σ*-path, viewed as a sequence of adjacent *σ*s, induces a unique ordered *η*-sequence, i.e., a 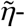 path in 𝒯 _*η*_(*t*). Together, these paths define an oriented trajectory 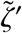, formed by uniquely directed straight links between adjacent locations, which can be graphically represented by a *directed polygonal chain* of *σ*-locations (Fig. 4A). Conversely, the fact that a generic 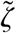 projects into a ordered sequence of adjacent *σ*-simplexes, 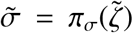, implies that oriented paths can be aligned into the polygonal chains, 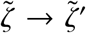 (Fig. 4B). From the perspective of the topological model discussed in Section II, this means that each path 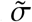 can be “lifted” from 𝒯 _*σ*_(*t*) into a unique oriented path 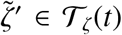 by a back projection, 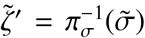. In the following, the term “simplicial path” will refer to the polygonal chains only, unless explicitly stated otherwise, and the “prime” notation will be suppressed.

**FIG. 4:**
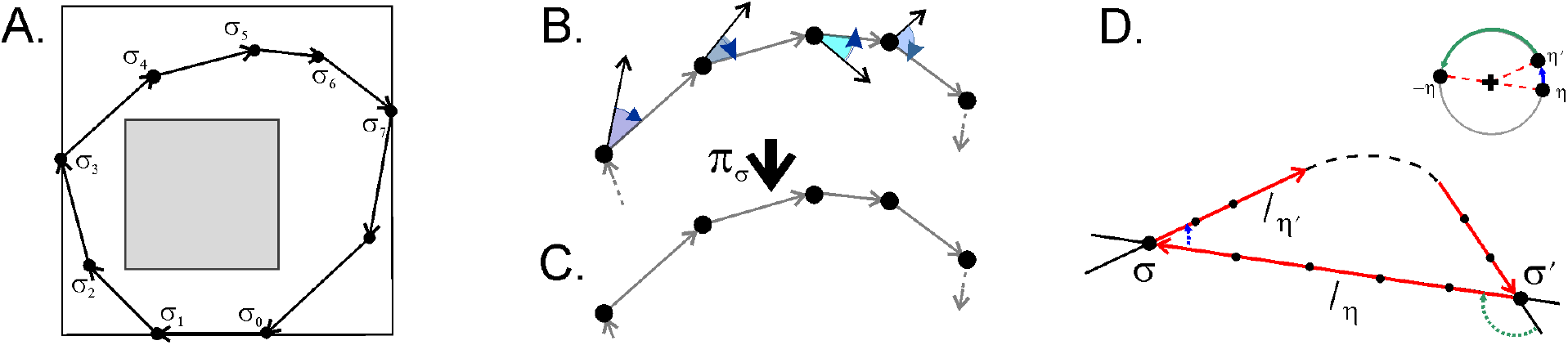
G-rules based geometric constructions. **A**. A directed polygonal chain connecting pairs of adjacent locations along the navigated trajectory in the environment shown on Fig. 1A. **B**. A schematic representation of a discrete homotopy from a generic *ζ*-path to a polygonal chain: the *η*-components shift towards the unique directions that align the adjacent locations (gray arrows). **C**. The resulting polygonal chain (a combination of yaws and straight runs) projects by *π*_*σ*_ into a undirected chain connecting the adjacent locations—a fragment of the chain shown on the panel **A. D**. Lemma 2: If two nonparallel alignments, 𝓁_*η*_ and 𝓁_*η*_′, produce two intersections *σ* and *σ*^*′*^, then there exists a 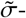 trajectory that forms a noncontractible simple loop, while the corresponding 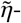 trajectory forms a contractible segment (top left corner), in contradiction with **R4**.

Reversing the order of simplexes in a chain and inverting the corresponding *η*-sequence,

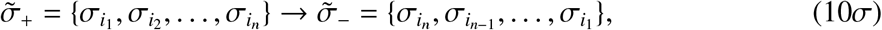

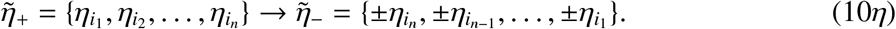

where the angle −*η* is diametrically opposite to *η, f*_*η*_(–*η*_*i*_) ≊ *π* + *f*_*η*_(*η*_*i*_). The “+” sign in (10*η*) corresponds to “backing up” along the path 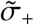 and the “–” sign to reversing the moving direction, either by implementing the required physical steps or by flipping the order of the replayed or preplayed sequences [81, 82] (in open fields, place cell spiking is omnidirectional [83]). The Since move reversal does not affect the *σ*-paths’ geometries, the transformations (10) can be regarded as equivalence relationships, which do not reference physical trajectory:

**R5. Reversibility**. *Simplicial σ-paths related via (10) are geometrically identical*.

In accordance with **R5**, a given *η*-oriented alignment,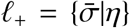, and its inverse, 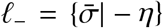, define the same collinear sequence, i.e., 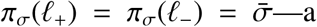 natural observation that motivates the definition **D7**. In particular, a pair of ±*η* adjacent locations is also geometrically adjacent 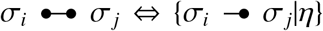 or 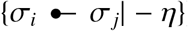, which allows representing trajectories by *undirected polygonal chains* connecting adjacent *σ*-fields (Fig. 4C). For example, a bending chain corresponds to a clockwise turn 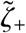 as well as to its counterclockwise counterpart 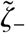 (both project to the same *σ*-path, 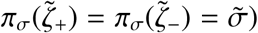; a closed chain—to a loop that can be traversed in clockwise or in counterclockwise direction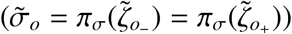, etc.

Returning to the link between the **G**-rules and the remaining two **A**- or **A**_*n*_-axioms, it can be observed that the **A2**_*n*_-axiom is an immediate consequence of **G2**: if the network is capable of actualizing up to *N*_*η*_ simplexes adjacent to a given one along *N*_*η*_ available directions, then *N*_*η*_ *−* 2 of them will necessarily lay outside of a given alignment. The argument for the existence and uniqueness of parallel lines (axiom **A3**_*n*_) can be organized into the following two lemmas:

### Lemma 1.

*If σ is a location outside of an alignment* 𝓁_*η*_, *σ* ∉. 𝓁_*η*_, *and* 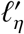*is an alignment directed by η at σ*, 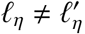, *then* 𝓁_*η*_ *and* 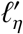 *do not overlap*.

**Proof**. Assume that the overlap exists, 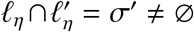. Since *η* is a unique index of directions, the location *σ*^*′*^ is *η*-aligned with its adjacent locations both in _*η*_ and in 𝓁_*η*_. Thus, 𝓁_*η*_ and 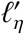 augment each other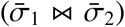, forming a single joint *η*-alignment that passes through *σ*, in contradiction with the original assumption *σ* ∉. 𝓁_*η*_. ■

### Lemma 2.

*Two non-parallel lines cannot intersect more than once*.

**Proof**. Consider two alignments 𝓁_*η*_ and 𝓁_*η′*_, *η≠ η*^*′*^, with 𝓁_*η*_ ∩ 𝓁_−*η*_ = *σ*. Without loss of generality (change 𝓁_*η*_ *→*_−_ 𝓁_*−η*_ if necessary), we may assume that the angle between them is sharp, ⦜(𝓁 _*η*_, 𝓁_*η*_) *< π/*2 (Fig. 4C). Consider an oriented path *ζ* that starts at *σ* in *η′* -direction, i.e., along _*η*_. If 𝓁_*η*_ crosses 𝓁_*η*_ again at a location *σ′*, then the path *ζ* may turn back at *σ* and continue along 𝓁_*η′*_ towards *σ*, then continue along 𝓁_*η′*_ again, etc., yielding a single closed *σ*-path. On the other hand, the corresponding *η*-path links *η* and *η′* at the first turn and then goes back from *η′* to −*η* at the second turn, forming a contractible segment, in contradiction with **R4**. ■

In effect, these two lemmas validate the constructive definition of parallelness **D7** and point at an alternative form of the axiom **A3**_*n*_: *If two locations σ*_1_ *and σ*_2_ *are aligned along η, then any two alignments* ℓ_1_ *and* ℓ_2_ *directed through σ*_1_ *and σ*_2_ *by a η*′ ≠ *η are parallel*.

The foregoing discussion suggests that the spatial framework represented by the place cells and the head direction cells forms neither a naïve discretization of the Euclidean plane nor a conventional finite geometry, as defined by the standard **A**-axioms. Rather, combining the location and the direction information can be used to capture a certain sub-collection of geometric arrangements, e.g., a particular set of alignments, which may then be used for navigation and geometric planning [20, 21]. Correspondingly, orientation learning can be interpreted as a process of establishing and expanding such arrangements (e.g., prolonging shorter alignments, completing partial ones, etc.) and accumulating them in the cognitive map.

## IV. SYNTHESIZING COGNITIVE GEOMETRY

As a basic example of a geometric map learning, consider an oriented trajectory *ζ*(*t*) that starts with an alignment 𝓁_1_ = {*σ*_0_, *σ*_1_ |*η*_1_ }, followed by a yaw at *σ*_1_, and continues along 𝓁_2_ ={ *σ*_1_, *σ*_2_ | *η*_2_ }, reaching *σ*_2_ at the moment *t*_2_ (Fig. 5A). As in Sec. II, the head and the motion directions are identified for simplicity. If *σ*_0_, *σ*_1_ and *σ*_2_ are the only locations in the emerging affine map *𝒜* (*t*_2_), then *σ*_2_ is adjacent to *σ*_0_ and hence it must align with *σ*_0_ along a certain *η*-direction *η*_20_ (assuming a generic case, in which 𝓁_1_ and 𝓁_2_ are nonparallel, *η*_1_ ≠ ±*η*_2_). Representing this alignment in the parahippocampal network, i.e., producing the corresponding imprints in the synaptic architecture via plasticity mechanisms [84–86], requires actualization by igniting *σ*_2_ and *σ*_0_ consecutively during the activity of a particular *η*_20_. This can be achieved either by navigating between the corresponding *σ*-fields or off-line, via autonomous network activity. In the former case, the connection 𝓁_20_ = {*σ*_2_, *σ*_0_ |*η*_20_ }is incorporated into the map after the animal arrives to *σ*_0_ from *σ*_2_, i.e., at the “empirical learning” timescale discussed in Section II. In the latter case, 𝓁_20_ may form at the spontaneous spiking activity timescale (milliseconds [18, 49–51, 59–62]), as soon as the animal reaches *σ*_2_, which clearly accelerates the formation of the cognitive map.

**FIG. 5:**
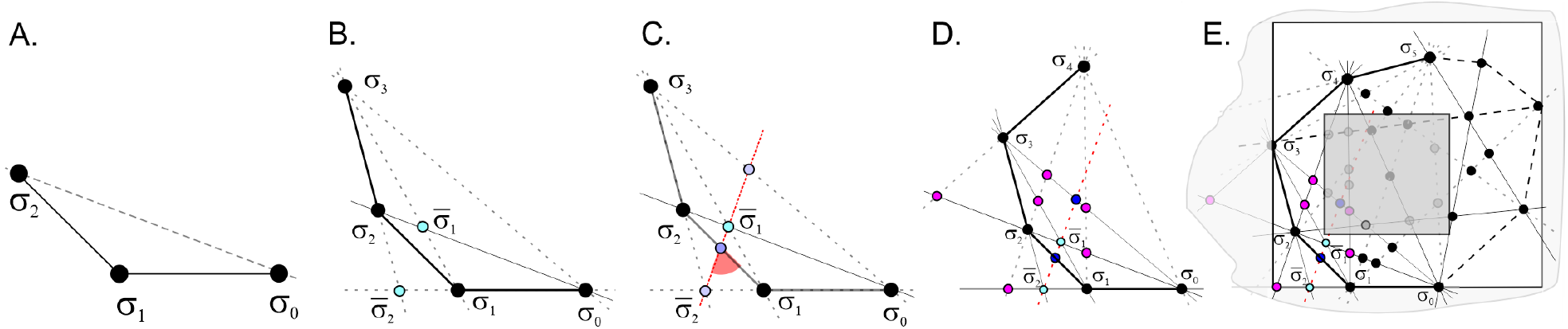
Aligning the cognitive map. **A**. The endpoints of the initial two segments of the trajectory are connected by a home run preplay 𝓁_20_ (dashed line). **B**. Reaching the next location, *σ*_3_, allows preplaying home runs to *σ*_0_ and *σ*_1_ and introducing the intersections 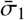 and 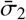 into the map. **C**. The direction of the alignment connecting the new points 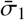 and 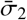 remain undefined, so 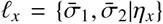 and may thus be viewed as provisional (topological) alignment in 𝒜 (*t*_3_). **D**. The location *σ*_4_ induces at several additional preplays to previously visited locations. **E**. At each step *t*_*n*_, the adjacent segments of the trajectory, 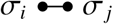 induce a *connectivity graph* 𝒢_*σ*_(*t*_*n*_), and the corresponding clique complex 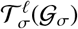 schematically represents the topological structure of the aligned map 𝒜(*t*_*n*_). **F**. The decays of the cell assemblies representing the unvisited locations (shaded areas) induces the required topological dynamics.

This illustrates the model’s general approach: although the geometric constructions were discussed in Section III in reference to spiking produced during the animal’s movements, they also apply to endogenous spiking activity. In other words, the **G**-rules can be used “imperatively,” for producing geometric structures in the cognitive map autonomously, based on available information rather than physical navigation. In particular, preplays can be used for aligning locations with specific *η*-assemblies by preplaying straight “home runs,” as soon as the physical trajectory assumes a suitable configuration.

The physiological processes that enforce transitions of activity between cell assemblies are currently studied both experimentally and theoretically [32, 33, 56]; in case of the head direction and place cells, the corresponding network computations may be guided by sensory (e.g., visual) and idiothetic (proprioceptive, vestibular and motor) inputs and involve a variety neurophysiological mechanisms [56, 87–90]. However, the principles of utilizing such mechanisms for acquiring a map of orientations can be illustrated using basic, self-contained algorithms that rely on the information provided by the hippocampal and head direction spiking. Specifically, the history of the *η*-assemblies’ ignitions allows estimating the direction between the loci of a polygonal chain according to

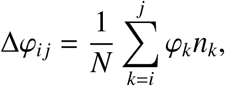

where *n*_*k*_ is the number of spikes produced by the *k*^th^ head direction assembly *η*_*k*_, *φ*_*k*_ is the corresponding angle (i.e., *η*_*k*_ = *π*_*η*_(*ζ*_*k*_) and *φ*_*k*_ = *f*_*η*_(*η*_*k*_)), *N* is the total number of spikes. If *η*_*k*_ is characterized by a Poisson firing rate *µ*_*k*_, then the number of spikes that it produces over an ignition period Δ*t*_*k*_ can be estimated as *n*_*k*_ = *µ*_*k*_Δ*t*_*k*_. Assuming for simplicity that all rates are the same *µ*_*k*_ = *µ*, the angular shifts can be estimated from the individual ignitions’ duration and the total navigation time *T*,

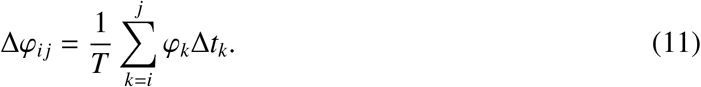

The *η*-simplex required to perform a home run from *σ*_*j*_ to *σ*_*i*_ can then be selected as the one whose discrete angle is closest to *φ*_*j*_ = *φ*_*i*_ + Δ*φ*_*i j*_, i.e.,

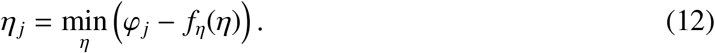

In particular, (11) and (12) allow estimating the required direction from *σ*_2_ to *σ*_0_ and thus identifying the simplex *η*_20_ that needs to direct the corresponding home run preplay 𝓁_20_. Other models can be built by modifying or altering these rules.

The next move continues along 𝓁_3_ = {*σ*_2_, *σ*_3_|*η*_3_}, arriving to *σ*_3_ at the moment *t*_3_, which allows preplaying connections to previously visited locations along 𝓁_31_ and 𝓁_30_ in the map *A*_C_(*t*_3_) (Fig. 5B). If *ζ*(*t*) is a right turn (*η*_1_ *< η*_2_ *< η*_3_), then the line 𝓁_31_ lays *between*𝓁 _30_ and 𝓁_3_, and, according to **G3**, overlaps with ℓ_20_ at 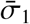, which will thus lay between *σ*_0_ and 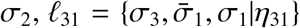.

Also, since 𝓁_3_ and 𝓁_1_ are non-parallel, they produce an overlap at 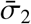 that extends the “seed alignments” 𝓁_1_(*t*_2_) and 𝓁_1_(*t*_3_) to 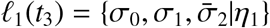 and 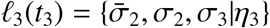. The locations within the alignments 𝓁_1_ and 𝓁_3_ are ordered correspondingly, e.g., *σ*_1_ falls between *σ*_0_ and 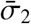, and *σ*_2_ falls between 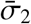 and *σ*_3_.

Note that since 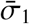 and 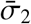 can be viewed as adjacent, it is also possible to form an additional alignment 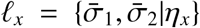, which induces two additional locations by intersecting 𝓁_30_ and 𝓁_12_ (Fig. 5C). However, the orientations of the existing segments of the trajectory do not determine the direction *η*_*x*_, and 𝓁_*x*_ can therefore be viewed as a “provisional” alignment that may be actualized once the explicit information specifying its orientation emerges. Nevertheless, such alignments and the incidences that they induce may be incorporated into the hippocampal map, to accelerate its topological dynamics.

As the turn continues, the next segment connects to *σ*_4_ along *l*_4_ = {*σ*_3_, *σ*_4_ |*η*_4_}, allowing home run preplays 𝓁_40_, 𝓁_41_, 𝓁 _42_, which produce additional intersections, augmenting the lines 𝓁_30_, 𝓁_31_ and 𝓁_43_ in specific order (Fig. 5D). Subsequent segments of the trajectory can generate ever larger sets of locations and alignments but the map learning process can be terminated when the map 𝒜_C_(*t*_*n*_) stabilizes topologically (see below). At each step, the acquired collection of alignments embedded into the unfolding cognitive map sustains its ongoing geometric structure.

Other alignments may be produced by more complex relationships within the existing configurations and their maps, as suggested, e.g., by Desargues or Pappus theorems. However, in finite planes these relationships may not be necessitated by the incidence axioms and require additional properties and implementing mechanisms [70–73]. This scope of questions falls beyond this discussion and will be addressed elsewhere.

### Topological quantification of geometric learning

The assumption of the model is that the influx of endogenously generated *σ*- and *ζ*-simplexes accelerates the emergence of an aligned cognitive map with the correct topological shape. Testing this hypothesis requires computing the persistent homologies of the corresponding *aligned complexes* 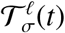 and 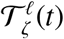 however, the algorithm described above produces the locations *σ*_*i*_ and their appearance times *t*_*i*_ without specifying cells that comprise a given simplex or detailing how these cells are shared between simplexes, which is required for the homological computations.

To extract the needed information, consider a graph 𝒢 _*σ*_ whose links correspond to the adjacent simplexes, i.e., vertexes *v*_*i*_, *v* _*j*_∈𝒢 _*σ*_ are connected if 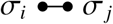. If each *σ*_*i*_ acts as an assembly, i.e., ignites when all of its vertex-cells (3) activate and if the adjacent simplexes share vertexes, i.e., 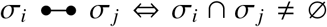 (required for spatiotemporal contiguity, see [74]), then each 𝒢 _*σ*_-link marks at least one putative cell *c*_*k*_ shared by *σ*_*i*_ and *σ*_*j*_. In a conservative estimate (assuming, e.g., no “redundant” cells that manifest themselves within just one assembly), the set of 𝒢 _*σ*_-links terminating at a given vertex *σ* thus defines the neuronal decomposition (3) of the corresponding simplex. Same analyses allow restoring neuronal decompositions for *η*-simplexes and constructing the *ζ*-simplexes, thus producing cliques simplicial complexes 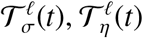 and 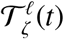.

If, according to the requirement **R1**, the resulting *σ*- and *η*-fields cover their respective representing spaces *ε* and *S* ^1^, then the *ζ*-fields cover the orientation space 𝒪 = *ε* × *S* ^1^, and the nerves associated with these covers, along with their temporal representations, 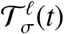 and 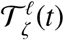, should have the required topological properties. However, this argument has a principal caveat: some locations induced through endogenous network activity may correspond to physically inaccessible domains in *ε*, which may divert the evolution of the resulting coactivity complex from the topology of the place field nerve of the navigated environment. Simulations show that indeed, the “autonomously constructed” complex 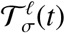 tends to acquire a trivial shape 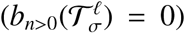 irrespective of the shape of the underlying *ε*.

A solution to this problem may be based on exploring functional differences between place cell combinations 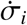 that represent “physically allowed” locations and the combinations 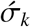 that represent “physically prohibited” regions. One would expect that in a confined environment, the former kind of cell groups should reactivate regularly due to animal’s (re)visits, whereas the latter kind is never “validated” through actual exploration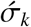s may activate only during the occasional preplays or replays. Taking advantage of this difference, let us assume that cell assemblies have a finite lifetimes [32, 33], i.e., that 1) the probability of an assembly’s disappearance after an inactivity period *t*_*σ*_ is

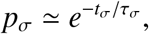

where *τ* _*σ*_ is *σ*’ mean decay period, and 2) that the decay process resets (*t*_*σ*_ = 0) after each reactivation of *σ* (for some physiological motivations and references see [67, 68]).

To emphasize the contribution of the locations imprinted into the network structure due to physical activity over the computationally induced locations, the latter may be attributed with a shorter decay period, 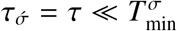, whereas the former may be treated as semi-stable 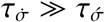 e.g., for basic estimates, one can use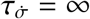. Lastly, the transition between 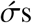and 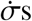 is modelled by stabilizing the decaying assemblies upon validation, i.e.,

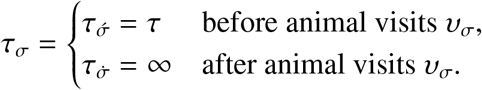

With this plasticity rule, physically permitted locations 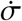should maintain their presence in the map, whereas the prohibited locations 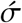 should decay, revealing the physical shape of the environment (Fig. 5E).

To verify this approach, the semi-random foraging trajectory simulated in Section II was replaced with a polygonal chain trajectory consisting of straight moves and random yaws. The preplays were then modeled by injecting straight alignments into the coactivity graph as soon as the required information became available (for details see [68]). Based on the results of [67, 68], the decay rate *τ* = 0.5 secs was selected to model the dynamics of the unstable locations.

For these parameters, the homological characteristics of the resulting “flickering” coactivity complex evaluated using ZigZag Homology techniques [91–93], quickly became stable: the Betti numbers stabilized at 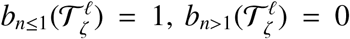 in 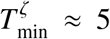 minutes, which approximately matches the hippocampal learning time 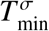 and demonstrates that geometric organization of the cognitive map brings learning dynamics to the biologically viable timescale.

## V. DISCUSSION

The proposed models of orientation learning are built by combining inputs from the hippocampal place cells and the head directions cells. Experiments demonstrate that these two populations of neurons are coupled: in slowly deforming environments, their spiking activities remain highly correlated, pointing at a unified cognitive spatial framework that involves both locations and orientations [27–30]. The goal of this study is to combine topological and geometric approaches to model such a framework, and to evaluate the corresponding learning dynamics.

The first0020model (Section II) is based on the observation that both the hippocampal and the head direction maps are of a topological nature: while the place cells encode a qualitative, elastic map of the navigated environment *ε* [13–18], the head direction cells map the space of directions, *S*^1^ [19]. A combination of place and head direction cells’ inputs can hence be used to construct an extended topological map of oriented locations *ο*, which has a structure of a direct product *ε* × *S*^1^—a natural framework for describing the kinematics of rats’ movements.

The second model (Section III) is structurally similar (a discrete map of directions is associated with each location), but involves constructions that define an additional, geometric layer of the cognitive map’s architecture. In particular, this model allows viewing spatial orientation learning from a geometric perspective—not only as a process of discovering connections between locations, but also establishing shapes of location arrangements, e.g., straight or turning paths, their incidences, intersections, junctions, etc. In neuroscience literature, such references are commonly made in relation to the physical geometry of the representing spaces *ε* and *S*^1^, e.g., the “straightness” of a *σ*-field arrangement implies simply that it can be matched by a Euclidean line in the environment where the rat is observed [20, 21]. However, understanding the geometric structure of the cognitive map requires interpreting neuronal activity in systems’ own terms, rather than through the parameters of exterior geometry.

The key observation underlying the geometric model is that the activity of head direction cells “tags” the activity of place cells in a way that allows an *intrinsic* geometric interpretation of the combined spiking patterns, i.e., defining alignments, turns, yaws, etc., in terms of neuronal spiking parameters. A famous quote attributed to D. Hilbert proclaims that “*the axioms of geometry would be just as valid if one replaced the undefined terms ‘point, line, and plane’ with ‘table, chair, and beer mug’*…” [94]. From such perspective, this model aims at constructing a synthetic “location and compass” neuro-geometry in terms of the temporal relationships between the spike trains, without using extrinsic references or *ad hoc* measures, which may be a general principle for how space and geometry emerge from neuronal activity.

In order to emphasize connections with conventional geometries, the model is formulated in a semi-axiomatic form. However, in contrast with the standard affine **A**-axioms or their direct analogues, the **A**_*n*_-axioms, the rules **G1**–**G3** serve not just as formal assertions that lay logical foundations for geometric deductions, but also as reflections of physiological properties of the networks that implement the computations. First, since the networks contain a finite number of neurons and may actualize a finite set of locations and alignments, the emergent geometry is finitary. Second, certain notions that in standard discrete affine plane 𝒜 are introduced indirectly, *relationally*, become *constructive* in the “cognitive” affine plane. For example, directions defined through equivalence classes of parallel lines in 𝒜 [70–73], are defined explicitly in 𝒜_*C*_ using the directing *η*-activities. Third, certain elements of the geometric structure are actualized explicitly through the network’s architecture, e.g., a fixed number of alignments passing through every location is implemented by cell assemblies wired into the head direction network [95, 96]. Other properties are not prewired but *acquired* during a particular learning experience and reflect both the physical structure of a specific environment and intrinsic mechanisms of spatial information processing.

## VI. METHODS

The computational algorithms used in this study were described in [6, 38–41, 67, 68, 74, 79].

**The environment** shown on Fig. 1A is similar to the arenas used in electrophysiological experiments [97, 98]. The simulated trajectory represents exploratory spatial behavior that does not favor one segment of the environment over another.

**Place cell spiking** probability was modeled as a Poisson process with the rate

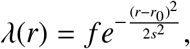

where *f* is the maximal rate and *s* defines the size of the firing field centered at *r*_0_ = (*x*_0_, *y*_0_) [99]. In addition, spiking probability was modulated by the *θ*-waves, which also define the temporal window *w* ≈250 ms (about two *θ*-periods) for detecting the place cell spiking coactivity [39, 100]. The place field centers *r*_0_ for each computed place field map were randomly and uniformly scattered over the environment.

**Persistent Homology Theory** computations were performed using Javaplex computational software [101] as described in [38–41, 79]. Usage of Zigzag persistent homology methods is described in [67, 68].

## Acknowledgments

The author is grateful to D. Morozov for providing Zigzag persistent homology simulating software. The work was supported by the NSF grant 1901338.

## Footnotes

1 Throughout the text, terminological definitions are given in *italics*.

